# Rule-governed Dynamic Stochastic Equilibration of Multicellular Motion *In Vivo* During Olfactory Neurogenesis

**DOI:** 10.1101/591479

**Authors:** Vijay Warrier, Celine Cluzeau, Bi-Chang Chen, Abigail Green-Saxena, Dani E. Bergey, Eric Betzig, Ankur Saxena

## Abstract

The complexity of patterning during organ-wide stem cell migration and differentiation can be challenging to interpret quantitatively. Here, we track neural crest (NC) and ectodermal placode-derived progenitor movements *in vivo*, for hundreds of cells, implement unbiased algorithmic approaches to extract biologically meaningful information, and discover cell-cell and lineage-lineage coordination between progenitors that form olfactory sensory neurons (OSNs) during zebrafish embryogenesis. Our approach discriminates between NC- and placode-derived contributions and segregates ingressing NC cells into two previously unidentified subtypes termed ‘trend’ and ‘dispersed’ lineages. Our analyses indicate that NC and placodal progenitor migration and intercalation are coordinated by at least two types of collective behavior: spatiotemporal exclusion and elastic tethering, akin to a push-pull mechanism. A stochastic equilibrium model accurately represents the interactions of NC and placode-derived lineages. Our approach provides insights into the coordination of dual-origin lineages during vertebrate olfactory neurogenesis and offers an algorithmic toolkit for probing multicellular coordination *in vivo*.

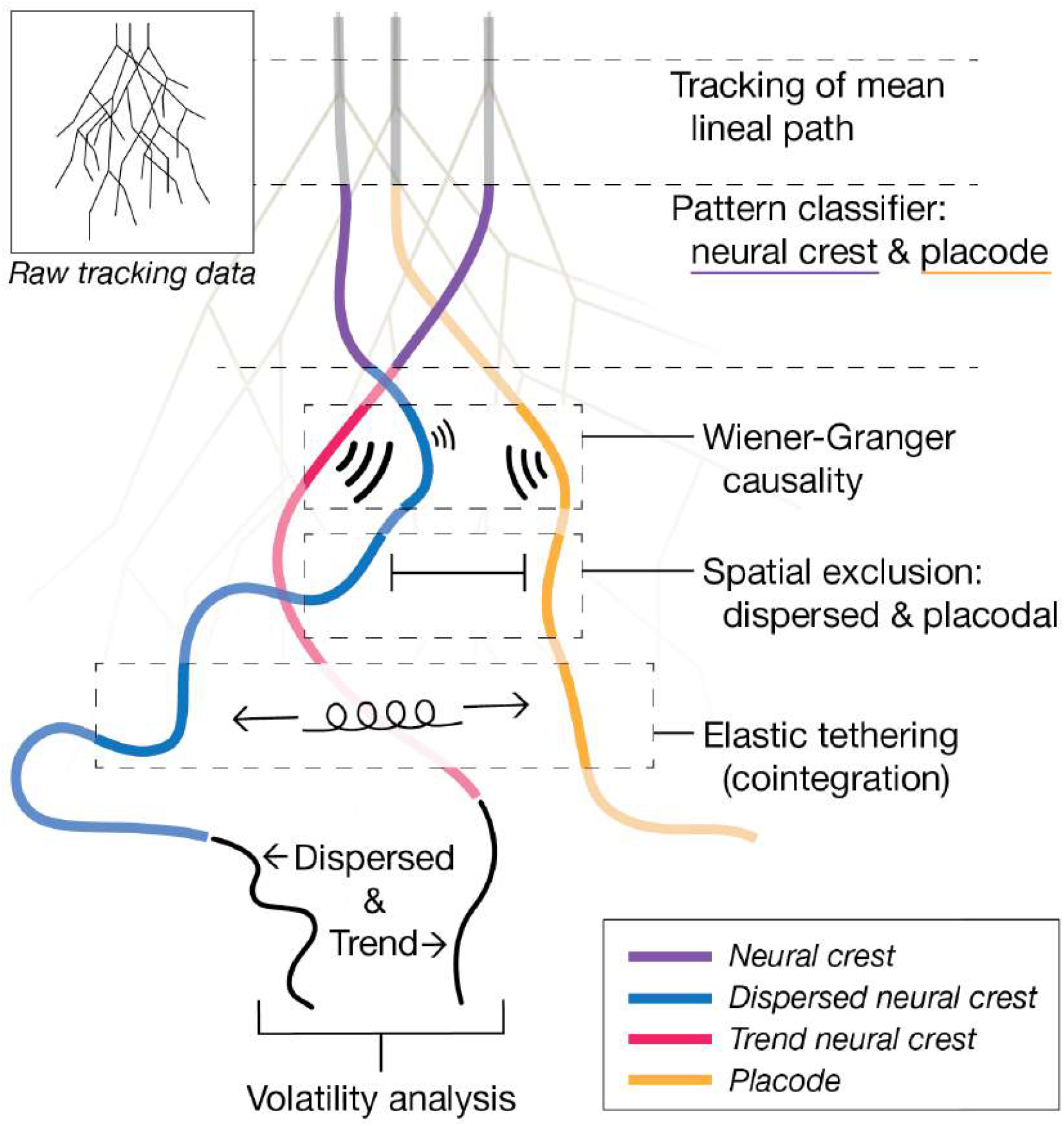
Graphical Abstract.

## Introduction

The multicellular dynamics of progenitor cell migration and differentiation are challenging to understand *in vivo*, particularly in densely packed organs such as the vertebrate olfactory epithelium (OE) where the system-level coordination of continuous, rapid neurogenesis remains poorly understood. Elucidating this process via phenotypic mapping at a multicellular and multilineage level is challenging, as disruptions to neurogenesis can cause system-wide phenotypes that, in turn, lead to secondary effects on tissue composition and/or organization. These primary and secondary effects can be difficult to separate out, in part due to a paucity of systems-level information on causal mechanisms across space and time. Recent advances in high-resolution imaging techniques, coupled with quantitative analysis, have the potential to provide a new window into *in vivo* interactions and causality during stem cell migration and differentiation.

The neural crest (NC) is a highly migratory stem cell population that traverses long distances to give rise to a variety of derivatives at multiple embryonic destinations (Dupin et al., 2018; Hall, 2018; Kuo and Erickson, 2010). A large body of work has described four broad categories of neural crest stem cells (NCCs) arranged along the anterior-posterior axis: cranial, cardiac/vagal, trunk, and sacral. Differences between these categories have been well-studied (Dupin et al., 2018; Hall, 2018; Kuo and Erickson, 2010), but little is known about behavioral variation within each category. Of note, pioneering work demonstrated that most, but not all, avian trunk NCCs synchronize cell cycle phase during emigration from the neural tube (Burstyn-Cohen and Kalcheim, 2002) and more recent findings showed significant differences in mitotic activity between subgroups of migrating cranial NCCs in chick embryos (Ridenour et al., 2014). While these cell cycle variations hint at a broader and complex underlying heterogeneity within at least some NC categories, the dynamic migratory behavior of subgroups remains unclear.

The most anteriorly-located NC population, the cranial NC, differentitates into many cell types that include cartilage, bone, neurons, and glia (Gallik et al., 2017; Rogers and Nie, 2018). Ectodermal thickenings termed placodes, meanwhile, give rise to major components of a number of sensory organs, including most of the OE (Baker and Bronner-Fraser, 2001; Maier et al., 2014; Moody and LaMantia, 2015). In zebrafish, the OE houses two major types of olfactory sensory neurons, ciliated and microvillous, with ciliated sensory neurons derived from the olfactory placode (Hansen and Zielinski, 2005; Hansen, 1993). The origins of microvillous sensory neurons, however, remain controversial. Our previous work showed that cranial NCCs migrate into the zebrafish OE to form the vast majority of microvillous sensory neurons (Saxena et al., 2013), while fate-mapping work from other groups across multiple species yielded both supportive (Forni et al., 2011; Katoh et al., 2011; Schott et al., 2018) and contradictory (Aguillon et al., 2018) findings, due, in part, to different assumptions about progenitor cell domains. Additionally, recent work suggests that neural plate border cells have greater than expected plasticity, with no clear demarcation between future placodal or NC cells (Roellig et al., 2017), adding further ambiguity to the debate. A shift in analysis away from spatial origin-based fate-mapping towards a holistic, multicellular categorization of differential behavioral characteristics may better illuminate these unique progenitor cell types and their descendants, but doing so requires both high-resolution data and unbiased quantitative methodology to tease out subtle cellular behaviors.

Traditional techniques such as confocal microscopy have limited utility for *in vivo* imaging due to difficult trade-offs between spatial resolution, temporal resolution, and phototoxicity, as well as resolution-based limitations on the accuracy and precision of automated data analysis. Therefore, to address the complexity of olfactory neurogenesis *in vivo*, we made use of lattice light-sheet microscopy (LLS; (Chen et al., 2014)), which balances exquisite temporal resolution with high lateral and axial spatial resolution without detectable phototoxicity or detrimental effects on growth. We imaged across time points that included continuous presumptive NC and placodal progenitor cell migration, intercalation, and differentiation in the developing zebrafish OE. LLS’s subcellular resolution allowed us to identify and track multiple cell lineages and opened up the potential to quantitate developmental processes that had been previously described only qualitatively. To overcome the challenges posed by large, complex datasets without introducing experimental bias inherent in the manual filtering of subsets of data, we developed unbiased modeling and analysis techniques that were inspired by the dynamic perspective on statistics found in econometrics for several decades (Granger, 1969). With this toolkit, we were able to capture high-density, multicellular dynamic patterning, i.e. motion over time, rather than the static patterning revealed by traditional fixing and staining protocols.

Previous quantitative analysis of olfactory neurogenesis has offered novel explanations of observed patterns of cell states and cell numbers based on a model of negative feedback dynamics within cell lineages (Lo et al., 2009). This study suggested plausible contributions to the stability of the OE by constructing a parametric model able to reproduce observed biological patterns and highlighted the importance of lineage-based analysis. Here, we endeavor to further understand lineage-dependent contributions to the OE by building models directly from empirical *in vivo* data that can then be statistically tested. Our approach preserves biological complexity by collecting and analyzing *in vivo* data at subcellular resolution that directly showcases the multicellular dynamics of progenitor cell migration during olfactory neurogenesis. We applied algorithmic pattern classification to high-resolution imaging data and, using cell lineage-based (lineal) trajectories, were able to effectively discriminate between progenitor cell types *in vivo*. Histogram and descriptive statistics suggested that lineal displacements are normally distributed. Adopting an interdisciplinary approach, we applied methods from financial portfolio theory to conduct a volatility analysis, which revealed that lineal displacements had a tendency to return to a trend value and identified two previously unrecognized subtypes of ingressing NCCs. These subtypes exhibited distinct behavioral characteristics in comparison to each other and to placode-derived cells, providing further fate-mapping independent evidence of an NC contribution to the OE. We next deployed the econometric method Wiener-Granger causality (WGC) (Granger, 1969), which provided statistical evidence that the tendency of displacements to return to a trend value could be modeled using a linear model. Finally, we compared the relative displacements of lineages with respect to one another to the well-studied problem of modeling stochastic deviation of asset price differentials about an equilibrium price spread (Johansen, 1991) and found strong statistical evidence that specific pairs of lineages are governed by a dynamic stochastic equilibrium. In doing so, we uncovered unexpected roles for specific cell lineages in guiding olfactory neurogenesis.

## Results

### High-resolution Tracking of Multicellular Dynamics During Olfactory Neurogenesis

The formation and morphogenesis of the OE, differentiation of olfactory sensory neurons, and outgrowth of their axonal projections to the olfactory bulb are carefully orchestrated developmental processes. We used LLS imaging to view these events at subcellular resolution. Figure 1A demonstrates laser scanning confocal and LLS images of the OE in live dual-transgenic zebrafish embryos displayed at the same resolution for comparison, with NC-derived microvillous neurons (Saxena et al., 2013) in green (*Tg(−4.9Sox10:eGFP*) (Carney et al., 2006; Wada et al., 2005)), referred to here as Sox10:eGFP, and proneuronal and neuronal nuclei in red (*Tg(−8.4neurog1:nRFP*) (Blader et al., 2003)), referred to here as Ngn1:nRFP. We applied LLS’s superior resolution in both space and time to follow the formation of the two main types of olfactory sensory neurons as they intercalated into the developing OE: microvillous (green, *Tg*(*TRPC24.5k:gap-Venus*)) and ciliated (red, *Tg*(*OMP2k:lyn-mRFP*)) (Sato et al., 2005) (Figure 1B, Video S1). Time-lapse sequences of up to 16 hours with temporal resolution of two to three minutes (2’-3’) (Figure 1B) exhibited no detectable phototoxicity or developmental delays during embryogenesis.

**Figure 1.**
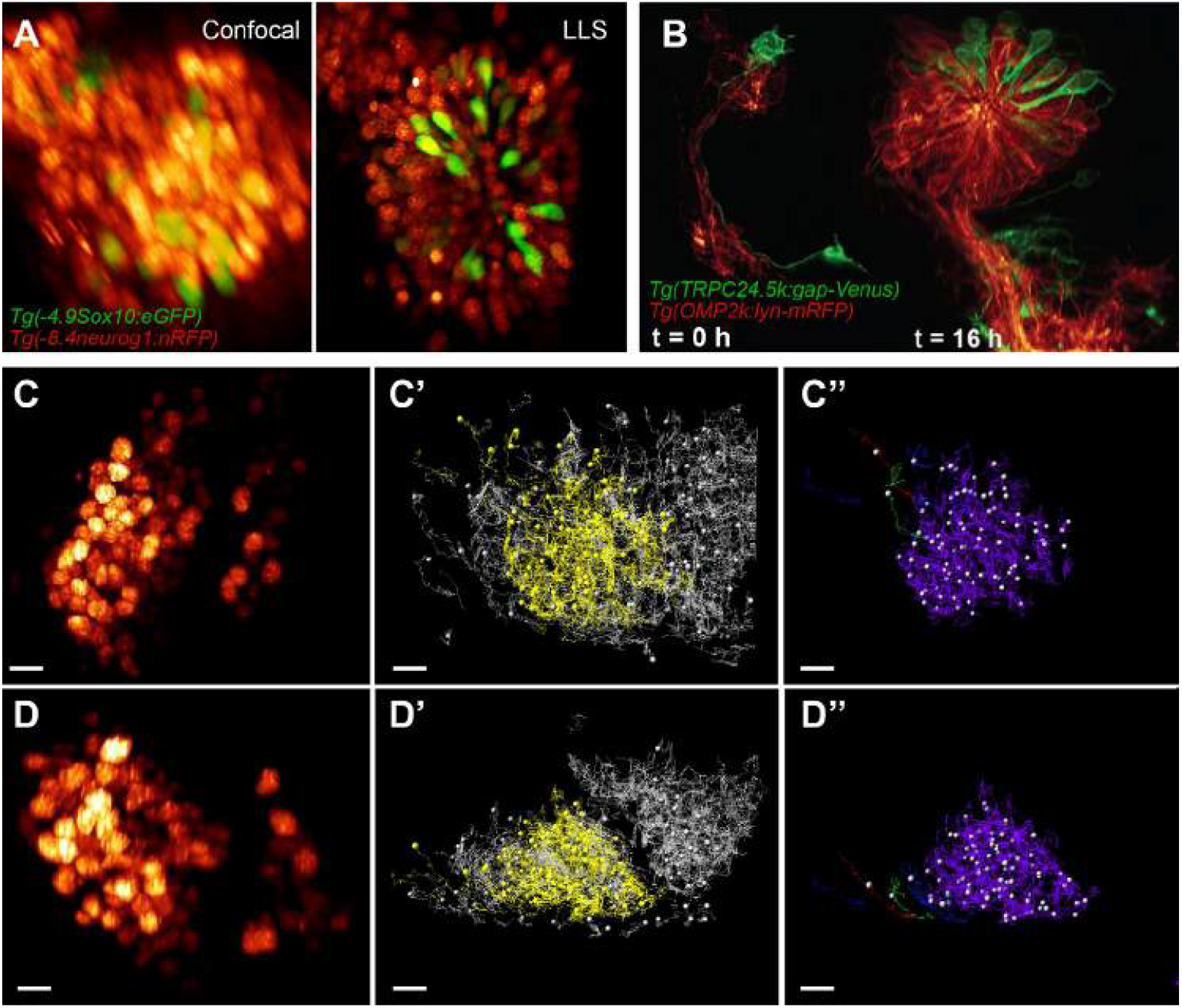
Imaging and tracking of progenitor cells and neurons in the developing zebrafish olfactory epithelium. (A) Representative examples of confocal and LLS imaging of the olfactory epithelium in live zebrafish embryos; cytoplasmic *Tg(−4.9Sox10:eGFP)* (green) marks cells with an NC origin and nuclear *Tg(−8.4neurog1:nRFP)* (red) marks neuronal fate specification. (B) LLS imaging of the developing olfactory epithelium over 16 hours yields visibly normal development of microvillous *(Tg(TRPC24.5k:gap-Venus)*, green) and ciliated *(Tg(OMP2k:lyn-mRFP)*, red) olfactory sensory neurons and their axonal projections to the olfactory bulb. (C-D”) Anterior (C-C”) and dorsal (D-D”) views of a *Tg(−8.4neurog1:nRFP)* dataset marking neuronal progenitors and neurons in the olfactory epithelium and bulb. (C’, D’) Migration tracks (grey) for all *Tg(−8.4neurog1:nRFP)*-positive cells at 24-30 hpf.; a subset of tracks are color-coded by displacement length in (C”, D”), with warmer colors representing longer displacements of putative NCCs. Scale bar: 10 μm.

LLS’s robust spatiotemporal resolution facilitated automated tracking of dense multicellular migration; Figure 1C-D” shows the migration tracks (Figure 1C’, D’) of several hundred Ngn1:nRFP-positive proneuronal or neuronal nuclei (Figure 1C, D) from 24 - 30 hours post-fertilization (hpf), with a subset sorted by displacement length (Figure 1C”, D”). During this time frame, we observed two broad categories of migratory behavior: 1) At the apical surface of the OE where neurons take residence, Sox10:eGFP-negative, placode-derived progenitor cells migrated relatively short distances (Figure 1C”, D”, purple); 2) Subsets of Sox10:eGFP-positive NCCs in the nasal cavity adjacent to the OE undertook longer migration paths (Figure 1C”, D”, blue, green, red) and ingressed into the OE to intercalate with placode-derived neurons. We next identified the spatial coordinates of every Ngn1:nRFP-expressing nucleus at 3’ time intervals in a dataset in which NC ingression into the OE at 24-29.5 hpf was clearly visible (Video S2) and constructed vectors for the direction and magnitude of each cell’s displacement. As cells divided and differentiated, we followed and annotated all observed cell lineages and constructed average lineal vectors for Sox10:eGFP-positive (NC) and Sox10:eGFP-negative (placodal) progenitors (Figure 2A). We limited our subsequent analysis to 63 NC-derived cells and 112 placode-derived cells that were clearly identified and grouped into 12 lineages and 20 lineages, respectively.

**Figure 2.**
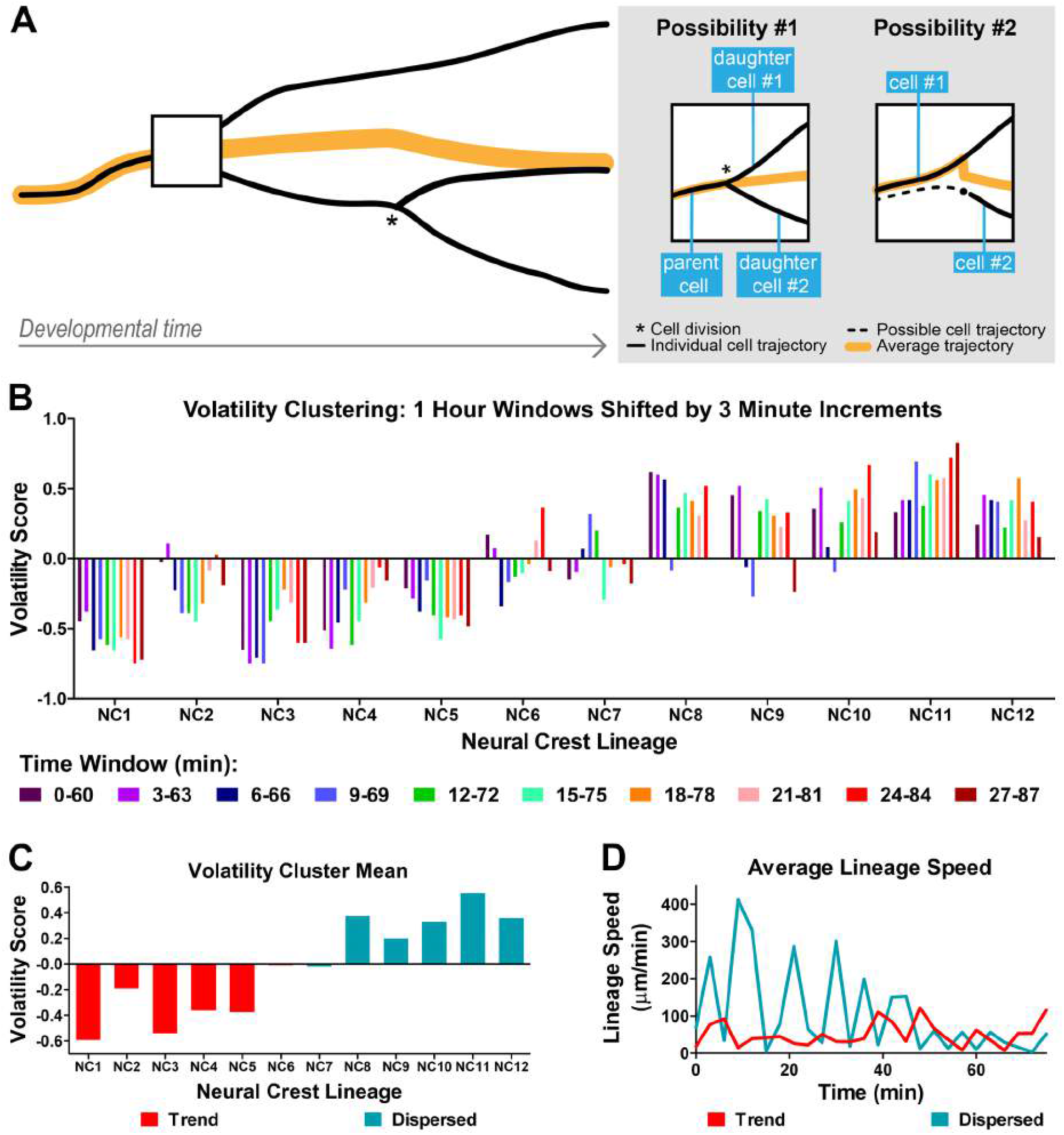
Volatility analysis of neural crest lineages with respect to average displacements identified two subpopulations of neural crest cells with differential migratory behavior. (A) Diagram outlining the trajectories of cells vs lineages. Individual cells were categorized into lineages using Imaris software’s ‘ImarisTrackLineage’ functionality. Lineal distributions were constructed from individual time series by averaging the positions of all cells from each lineage at each time point. (B) Volatility scores for each NC lineage, calculated for 10 consecutive 60’ time windows, shifted by 3’ increments, at 24-25.5 hpf. Average lineal positions for every pair of NC lineages (12 lineages, 66 pairs) were calculated. The volatility score (square root of mean squared displacement from pairwise trend) was calculated for each lineage. Volatility clustering appears robust to time-window shifting. (C) Sum of volatility scored values for each NC lineage demonstrating a distinctive bias towards negative scored values, indicating low volatility, for trend lineages and a distinctive bias towards positive scored values, indicating high volatility, for dispersed lineages. (D) Average speed of the two clusters of neural crest lineages demonstrating the significantly lower average speed of trend lineages compared to dispersed lineages.

### Algorithmic Training Using Lineal Tracks Yields Accurate Classification of Cell Types and Origins

We first analyzed cellular displacements on all three embryonic axes: anteroposterior, dorsoventral, and lateromedial. We observed no spatial bias in the starting position of ingressing NCCs, and a principal components analysis indicated that 70%,16%, and 14% of the net displacement for NC-derived cells occurred along the anteroposterior, dorsoventral, and lateromedial axes, respectively, as compared to 40%, 23%, and 37% for placode-derived cells. These results are consistent with the visually-established, predominantly anteroposterior motion of the cranial NC at these developmental stages (Saxena et al., 2013) and the more broadly distributed displacements of placodal derivatives.

We next asked whether quantitative characteristics of Ngn1:nRFP-positive cells’ motion could reliably distinguish between NC- and placode-derived cell populations independent of fluorescent marker specificity. Starting with our total Ngn1:nRFP-positive, dual-origin cell population, we manually selected the subset of cells that was Sox10:eGFP-positive and used these data to train an algorithmic cell classifier based on several combinations of cellular trajectory statistics (see Methods for details). However, every combination implemented correctly classified only 50% or less of cells (data not shown), a threshold not sufficient to be analytically useful. Therefore, we next attempted training with lineage trajectory statistics and found a dramatic improvement in classification. Our most robust automated classifier, based on anteroposterior net displacement, anteroposterior velocity, and dorsoventral net displacement, correctly categorized 87.5% (28/32) of lineages (both NC and placode) when benchmarked against our fluorescent-marker based identities. Thus, lineage-based tracks proved highly informative for the purposes of identifying NC or placode derivation based solely on spatiotemporal movement characteristics.

### Volatility Analysis Reveals Dynamically Distinct Subtypes of Neural Crest-Derived Lineages

NCCs differentiate into a large number of cell types, and in the OE and surrounding nasal cavity, they give rise to sensory neurons and cartilage precursors, respectively (Dale and Topczewski, 2011; Saxena et al., 2013). Little is known about how and when the specification of these cell types occurs, but slight differences in cell positions and/or microenvironments could potentially influence NC behavior. Therefore, we performed a volatility analysis to test whether NC-derived lineages demonstrated heterogeneity in their behavior, wherein volatility is a measure of the dispersion of a displacement time series centered on a displacement trend value rather than the mean displacement. We constructed a novel approach to calculate volatility without making assumptions about the myriad molecular mechanisms or spatiotemporal distribution of signals that cells could be responding to and focused this analysis on a 90’ time span of data (24-25.5 hpf) when ingression of NCCs into the OE was rapid, frequent, and clearly visible. We defined displacement trends for each of the 12 NC-derived lineages as the net displacement over this 90’ window, calculated the volatility of the displacements around these trends, then calculated the volatility of each lineage with respect to the trends of other lineages (66 total pairings of trends with lineages), and devised a volatility score normalized to lie between −1 and 1, with negative scores representing greater adherence to a trend and positive scores representing a greater deviance, i.e. ‘dispersed’ behavior (Figure 2B-C). In order to rule out the possibility that a specific time window could present artifactual volatility, we calculated volatility scores over 10 overlapping time windows (Figure 2B). Strikingly, NC-derived lineages segregated into two distinct clusters (Figure 2C): a ‘trend’ cluster with low volatility (lineages 1-5) and an average normalized score over all time windows of −0.42; and a ‘dispersed’ cluster with high volatility (lineages 8-12) and an average normalized score over all time windows of 0.37. Lineages 6 and 7 had intermediate volatility scores and could not be classified based on clustering alone.

Further analysis of the two clusters revealed remarkably distinct properties of motion, with trend lineages exhibiting a far lower velocity than did dispersed lineages. Including these parameters allowed us to classify lineages 6 and 7 in the trend and dispersed clusters, respectively. The velocities for lineages 1-6 (trend) yielded an average value of 50.14+/−25.61 μm/min whereas those of lineages 7-12 (dispersed) yielded a significantly different average value of 104.45 +/− 92.95 μm/min (p<0.02). Both the trend and dispersed populations exhibited oscillating variation in their velocities, and the dispersed cluster exhibited rapid speed fluctuations (as reflected in its wide standard deviation) that were not seen in the trend cluster (Figure 2D). In sum, the initial volatility analysis coupled with subsequent analyses of motion has illuminated previously unidentified subtypes of NCCs that ingress into the developing OE.

### Spatial Exclusion Exists Between Placode-Derived and Dispersed Neural Crest-Derived Lineages

In addition to marked differences in volatility and rate of movement, we also identified differential spatial characteristics for trend and dispersed NC lineages. Over the full time-course of our dataset, dispersed lineages were spatially clustered around the lumen of the OE (Figure 3A-D; Video S3), whereas trend lineages were anteriorly concentrated and diffused across the dorsoventral and lateromedial axes (Figure S1; Video S4). Intriguingly, dispersed lineages, in contrast to trend lineages, presented no spatiotemporal overlap with placode-derived lineages, i.e. the full analyzed time frame of migration had zero intersecting dispersed and placode-derived trajectories across both space and time (Figure 3A-D; contrast with trend lineages in Figure S1; Videos S3 and S4). These spatial exclusion data and the volatility analysis above, taken together, provide the first identification of two distinct subtypes of cranial NC-derived lineages and describe their unique behaviors and migration patterns during olfactory neurogenesis.

**Figure 3.**
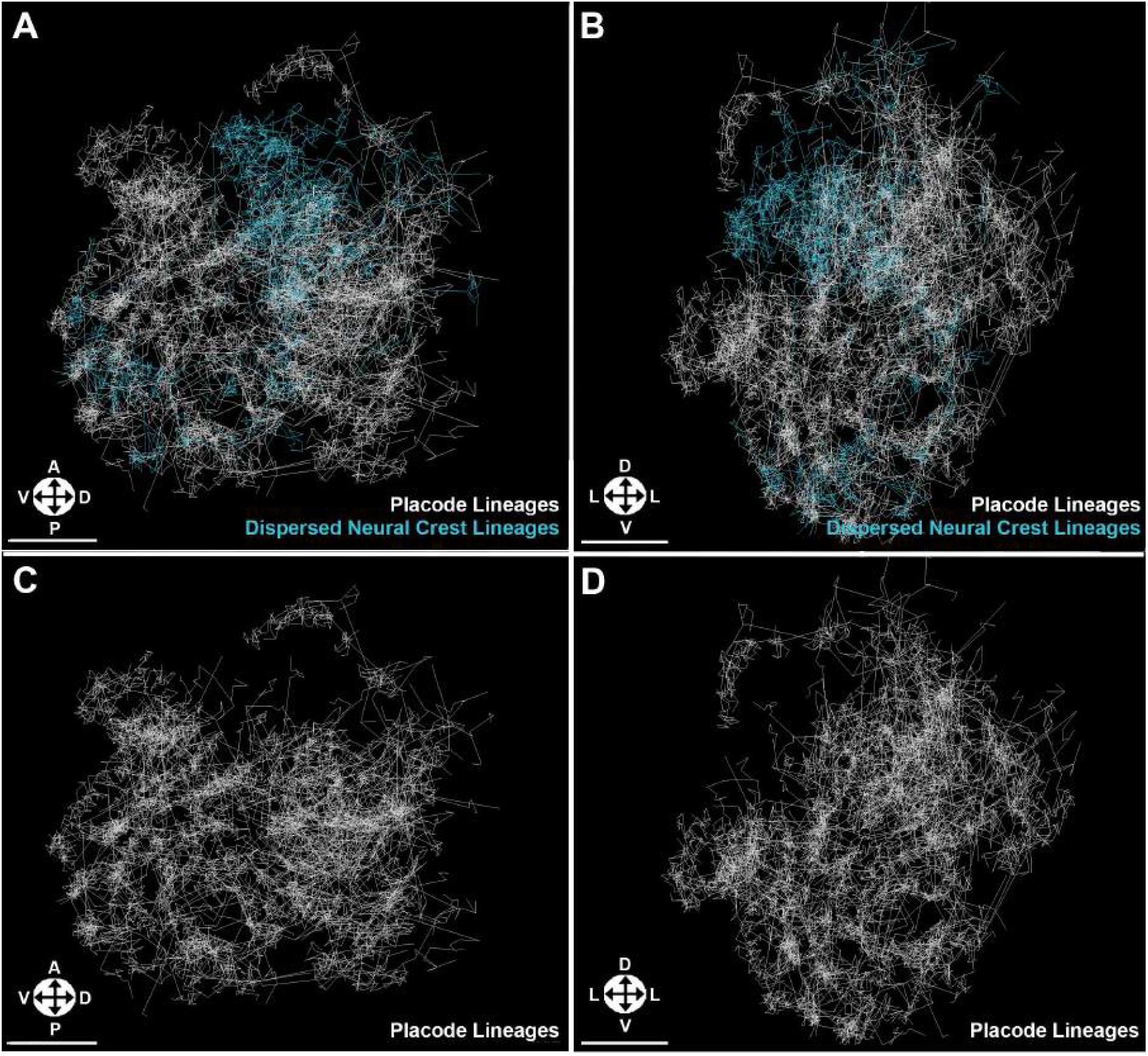
Spatial Exclusion of Dispersed Neural Crest Lineages and Placode Lineages. (A, C) Lateral views of the olfactory epithelium showing trajectories of individual dispersed NC (blue) and placodal (white) lineages at 24-29.5 hpf. (B, D) 90 degree rotations of views shown in (A, C). Dispersed NC and placode trajectories do not overlap during the full 5.5 hours time span examined, whereas trend trajectories (Figure S1) do. Orientation arrows: A, anterior; P, posterior; D, dorsal; V, ventral; L, lateral. Scale bars: 15 μm.

### Causal Analysis of Cellular and Lineal Trajectories Indicates Directed Information Between Lineage Types

Moving beyond observational analysis, we wanted to determine if our novel identification of lineage subtypes and their quantitative characteristics would allow for unbiased predictions of behaviors and interactions. We first tested individual cell tracks for WGC, which provides statistical evidence of information-theoretic predictive relations between time series (Granger, 1969). In brief, a statistically significant finding of WGC between trajectories X and Y suggests that a prediction of Y’s position based on Y’s past values is improved by including X’s past values. As a whole, our WGC results for individual cell trajectories exhibited levels of statistical significance that spanned many orders of magnitude (Figure 4A), which made them difficult to interpret. Of interest, however, were specific trajectories with the most statistically significant WGC (p<0.001) that, upon visual comparison, were found to demonstrate similar patterns of motion in all four dimensions (Figure 4B). These results pointed to a limited but promising utility for WGC analysis on its own, but the ubiquity of statistically significant (p<0.05) WGC relations suggested that a more refined approach was needed in order to efficiently discern pattern relations between a large number of trajectories.

**Figure 4.**
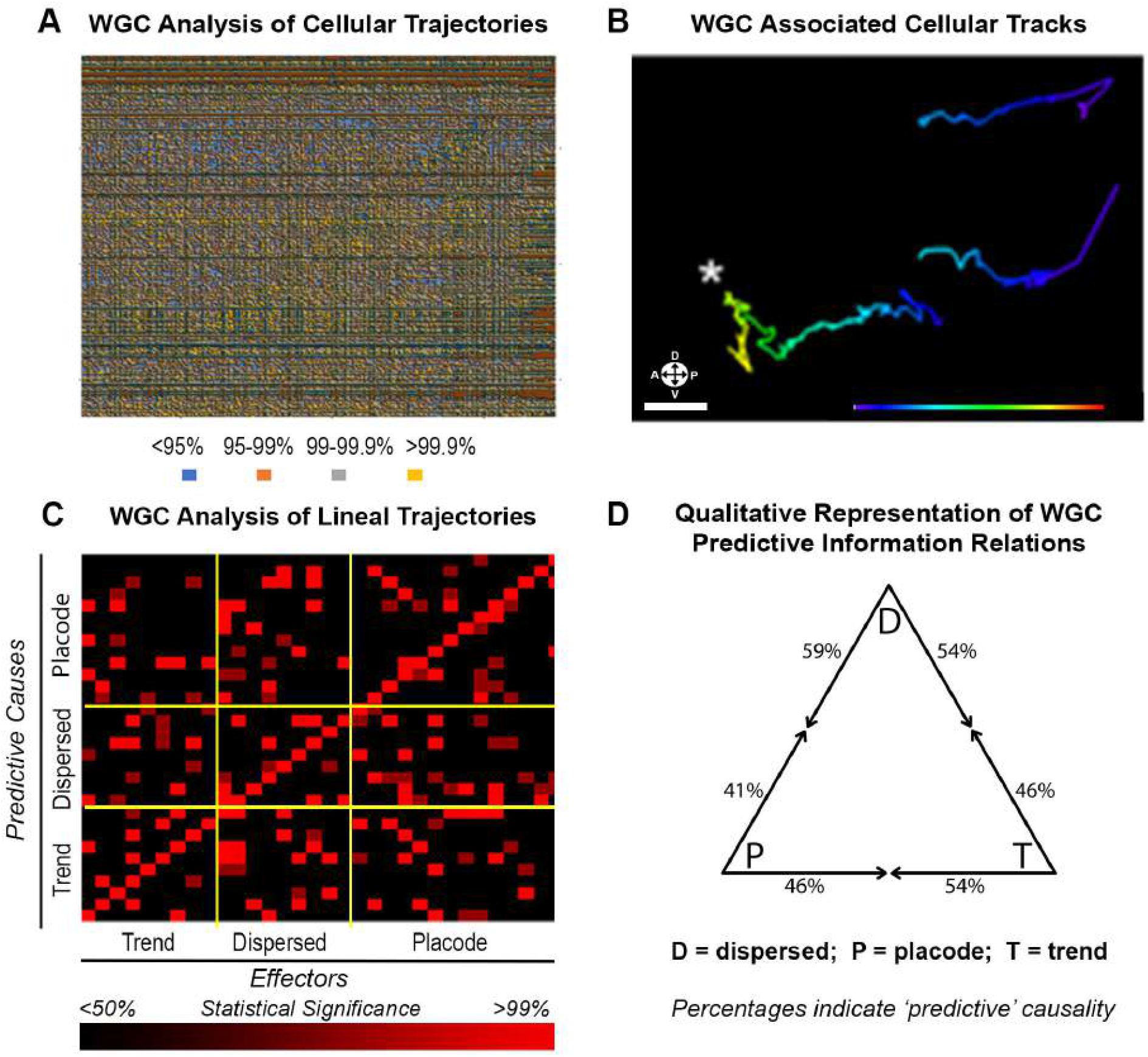
WGC Analysis Facilitates Selection of Qualitatively Similar Cell Tracks and Provides Evidence of Direction of Predictive Information Flow Between Lineages. (A) Heat map indicating WGC relations between 250 cellular trajectories analyzed pairwise at 24-25.5 hpf, color-coded by level of statistical significance. (B) Selected cell tracks related by WGC causality at >99% statistical significance; asterisk indicates the WGC ‘predictively causal’ track, while the other tracks are ‘effectors’ with similar spatiotemporal migration profiles. (C) Heatmap indicating WGC relations between all analyzed lineal trajectories in dataset at 24-25.5 hpf, color-coded by level of statistical significance. (D) Apparent directed flow of predictive information as quantitatively evidenced by the heatmap in (C). Predictive information from trend and dispersed lineages to placode lineages is not symmetric; instead, more predictive information flows from the neural crest lineages to the placode as compared to the reverse flow. Orientation arrows: A, anterior; P, posterior; D, dorsal; V, ventral. Scale bar: 10 μm.

Given the previously described improvement in algorithmic discrimination between NC- and placode-derived identities when shifting from cellular to lineage trajectories, we considered WGC relations between lineages (Figure 4C). This approach greatly reduced the fraction of WGC relations of statistical significance exceeding 95%. Restricting our attention to these statistically significant relations and taking advantage of the directed character of WGC relations (if time series X significantly predicts time series Y, time series Y need not predict Y), we calculated the asymmetry of WGC relations (Figure 4D) along the anteroposterior, dorsoventral, and lateromedial axes. These calculations provide an overview of the relative balance of predictive information flow along each analyzed axis. NC-derived dispersed lineages exhibited greater predictive information flow in relation to placode lineages at a ratio of 13:9; NC-derived trend lineages exhibited lesser predictive information flow in relation to dispersed lineages at a ratio of 6:7; NC-derived trend lineages exhibited predictive relation flow in relation to placode lineages at a ratio of 7:6. These findings suggest that the mechanisms responsible for the observed predictive relations of lineal motion may act in asymmetric ways on the different types of lineages and/or that the distinct properties of lineages generate asymmetric correlations from uniform mechanisms.

### Cointegration Reveals a Dynamic Stochastic Equilibrium of Lineal Displacements

We next endeavored to fit the statistical dependencies revealed by WGC to a dynamical model that could help explain the coordinated migration and rearrangement of NC- and placode-derived cells. Given the sinusoidal patterning of lineal speeds (Figure S2A) and normal distributions of lineal displacements (Figure S2B), we applied cointegration, a variant of WGC analysis that statistically tests a model of dynamical equilibration based on a harmonic oscillator mechanism subject to normally-distributed perturbations. The finding of cointegration between two stochastic time series can be understood on analogy to ‘elastic tethering’: the time series will vary over time, but differences between cointegrated series tend to equilibrate around a mean separation distance, as if they are elastically tethered.

Since principal components analysis had indicated that NC migration during olfactory development is primarily anterior-posterior, we performed a cointegration test on the anterior-posterior component of cellular motion. We calculated cointegration relations between three types of trajectories: 1) nonmigrating, cartilage-fated NC cell trajectories in the nasal cavity as a negative control (n=10, Figure 5A, ‘Stationary NC Cells (Control)’); 2) migrating placode- or NC-derived cell trajectories (n=66, Figure 5A, ‘Placode and Mig. NCCs’); 3) placode- or NC-derived lineage trajectories (n=32, Figure 5A, ‘Placode and Mig. NC Lineages’). Importantly, there was no statistically significant cointegration of non-migrating, cartilage-fated NCCs in the nasal cavity with respect to the other trajectories (Figure 5A), as would be expected for this stationary control group of cells. We also found an absence of cointegration relations for migrating lineages in the nearby but separate olfactory bulb (data not shown), suggesting that our test is not producing non-specific cointegration. We next looked at individual cell trajectories and observed almost no statistically significant cointegration of time series (Figure 5A). Lineage trajectories, on the other hand, evidenced 49 pairs of lineages with statistically significant cointegration (p<0.05; Figure 5A, B). These results suggest that the mechanisms responsible for elastic tethering act with lineal specificity. Intriguingly, the cointegration relations exhibited by the previously identified trend and dispersed NC-derived lineages were highly distinct (Figure 5B, C). Trend lineages did not reach the threshold of 95% statistical significance for cointegration among themselves and were not significantly cointegrated with respect to dispersed and placode lineages. By contrast, multiple dispersed lineages were significantly cointegrated with each other and with placode lineages (Figure 5B, C).

**Figure 5.**
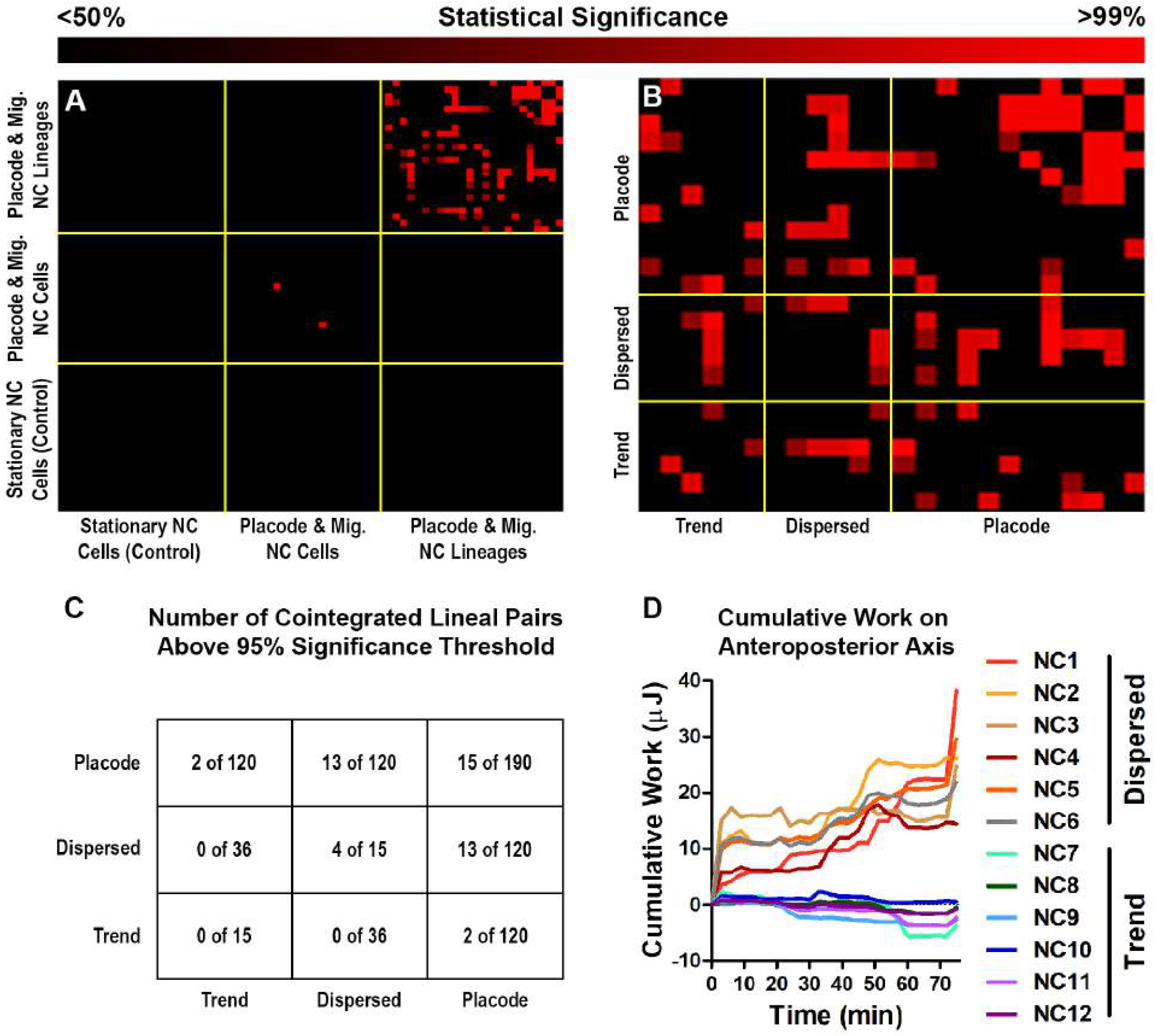
Cointegration enables quantitative identification of patterns of statistical dependency between lineal trajectories. (A) Heat map of the outcome of cointegration tests. Red/black coloring corresponds to higher/lower values of the F-statistic generated by the cointegration test, color-coded by statistical significance. Rows and columns represent trajectories at 24-25.5 hpf for Stationary NC cells (Control; all null) in the nasal cavity; placode and migratory NC cells; or placode and migratory NC lineages. The top right region of (A) is expanded in (B) to demonstrate the F-statistic for each pair of analyzed lineages segregated by lineage type. (C) Trend NC lineages are not cointegrated at >95% with any NC lineages and are cointegrated at >95% with 2 placode lineages, whereas dispersed NC lineages are cointegrated with 4 dispersed NC lineages and 13 placode lineages at >95%. Additionally, placode lineages are cointegrated at >95% with 15 placode lineages. (D) Cumulative work (in units proportional to micro-joules) for each NC lineage done along the anterior-posterior axis by the sum of all the elastic tethers implied by the cointegration relations over 72’. Trend NC lineages demonstrate a minimal accumulation of work (mean across all trend lineages at final time point = −1.7 +/− 1.2 ∝:μJ, p = 0.05), whereas dispersed NC lineages demonstrate a progressive accumulation of work (mean across all dispersed lineages at final time point = 25.7 +/− 6.4 ∝:μJ, p = 0.05). Cointegration half-life analysis confirms that trend and dispersed NC lineages exhibit differential migratory behavior.

Based on our elastic tethering model, we calculated an elasticity coefficient for each lineage representing how deviations from the equilibrium separation between lineages are corrected. Taking into account the calculated distance between lineages at each time point, these elasticity coefficients enabled us to estimate the force exerted by statistically significant tethers on each individual lineage. These approximations were used to generate cumulative work estimates associated with the directed movement of NC-derived lineages, producing values ranging from 0 to 40 units proportional to μJ (Figure 5D). A significant (p<0.0001) distinction between trend and dispersed NC lineages emerged, with trend lineages demonstrating a minimal accumulation of work (mean at final time point = −1.7 +/− 1.2 units proportional to μJ) and dispersed lineages demonstrating a progressive accumulation of work (mean at final time point = 25.7 +/− 6.4 units proportional to μJ). These findings are consistent with our visual observations of steadily progressing ingression of NC lineages into the OE along the anteroposterior axis (Video S2).

## Discussion

Here, we used high-spatiotemporal resolution imaging to view migratory behaviors of distinct cell populations in the zebrafish OE at a level of detail not previously possible, which in turn allowed us to accurately identify and track the movements of individual cells and their resulting cell lineages. Our analyses, which did not require *in silico* modeling but rather were built directly off of *in vivo* observations, yielded previously unknown patterns of progenitor multicellular movements, the categorization of newly identified subtypes of NCCs, and the identification of causal relationships between specific cells and lineages. Previous studies employing advanced imaging modalities have most often taken advantage of superior resolution to draw qualitative conclusions (Wang et al., 2014) or study quantitative phenotypes that were previously not discernible using lower-resolution modalities (Aguet et al., 2016). Our approach differs in that we have used the superior resolution of LLS to directly input into and enable statistical testing of a dynamical model.

Pursuant to this goal, we employed multiple quantitative approaches that were inspired, in large part, by longstanding innovations in the study of econometrics and financial theory. One approach, WGC, has been previously applied to better understand nervous system function and activity (Bressler and Seth, 2011) but not, to our knowledge, to study nervous system development. Our multipronged strategy included algorithmic cell and lineage type discrimination based on statistical pattern classification, mean-variance volatility analysis, WGC, and cointegration, cumulatively yielding new insights into progenitor cell dynamics during olfactory neurogenesis. These analyses produced several noteworthy quantitative findings: dynamical information is concentrated in lineal positions and velocities rather than in individual cell data; there are two dynamically distinct subtypes of ingressing NCCs; the predictive relations between the placode lineages and the two types of NCC lineages are quantitatively non-equivalent; and there is strong statistical evidence that NC ingression stochastically equilibrates about a mean trend of motion.

The improvement in performance of our automated cell classifier, WGC analysis, and cointegration tests upon using lineage data suggests that cell lineage behavior is more informative than individual cellular data of *in vivo* multicellular behavior during olfactory development. This supposition is consistent with the idea that the quantitative fingerprint of any single cell-level mechanism is likely to be obscured by factors such as the heterogeneity of cellular microenvironments and spatial variability in individual cells’ responsiveness to signals. Thus, the aggregate measurements provided by lineage analysis can illuminate collective cell dynamics more robustly than can individual cell measurements, analogous to how econometrics can illuminate aggregate economic quantities.

The two new subgroups of cranial NC that we identified, trend and dispersed, behave differently with respect to placode-derived cells as evidenced by spatial exclusion, causal relations, and cointegration/elastic tethering analyses. The curious observation of spatiotemporal exclusion between dispersed and placode-derived trajectories (in contrast to that between trend- and placode-derived trajectories) suggests an additional level of complexity in the coordination of NC ingression and placode rearrangement. Our observations of statistically significant cointegration relations between time series of lineal motion provide strong statistical support for the claim that NC ingression exhibits a dynamic stochastic equilibration (elastic tethering) around a trend of motion and offer indirect, fate-mapping independent evidence of an NC contribution to OSNs.

Given the stochastic nature of embryonic development, single-cell comparisons *in vivo* are difficult to make, whereas comparative multicellular behavior, analogous to viewing a large wave rather than a single ripple, can move forward our understanding of system-wide biology. The results presented here, obtained via unbiased mathematical analyses, reveal a highly dynamic interplay among different progenitor cell types during the formation of the olfactory system. This multilineage coordination and causality had remained undetected using traditional methodologies, and our findings of dynamical distinctions between ingressing NC lineages and of new behavioral phenomena were made possible by converting high-spatiotemporal resolution imaging into numerical values that were fed into complex algorithms deployed to advance an understanding of system-wide *in vivo* data. Moving forward, this general approach offers an algorithmic toolkit for probing multicellular coordination and, if applied to genetic perturbations, may help elucidate causal mechanisms *in vivo* via the discrimination of subtle spatiotemporal phenotypes.

## Acknowledgements

We thank Dr. Robert Kelsh for the Sox10:eGFP line; Dr. Uwe Strahle for the Ngn1:nRFP line; Dr. Yoshihiro Yoshihara, RIKEN BSI, and the National Bioresource Project of Japan for the OMP:RFP and TRPC2:Venus lines; the HHMI Janelia Research Campus Vivarium Staff for zebrafish care; Dr. Marianne Bronner for generously supporting the initiation of this project and for valuable feedback on the manuscript; and Mr. Govind Warrier for insightful conversations regarding analytical methodology. This work was funded by the Chicago Biomedical Consortium with support from the Searle Funds at The Chicago Community Trust, an HHMI Janelia Visiting Scientist Program Research Support Award, and University of Illinois Lab Startup Funds.

## Author Contributions

V.W., B-C.C., E.B., and A.S. conceived and designed all experiments and analysis. V.W., B-C.C., and A.S. performed and analyzed all experiments. V.W., C.C., A.G-S., D.E.B., and A.S. designed and made figures. V.W., C.C., A.G-S., and A.S. wrote the manuscript.

## Declaration of Interests

The authors declare no competing financial interests.

## Supplementary Material

**Figure S1.**
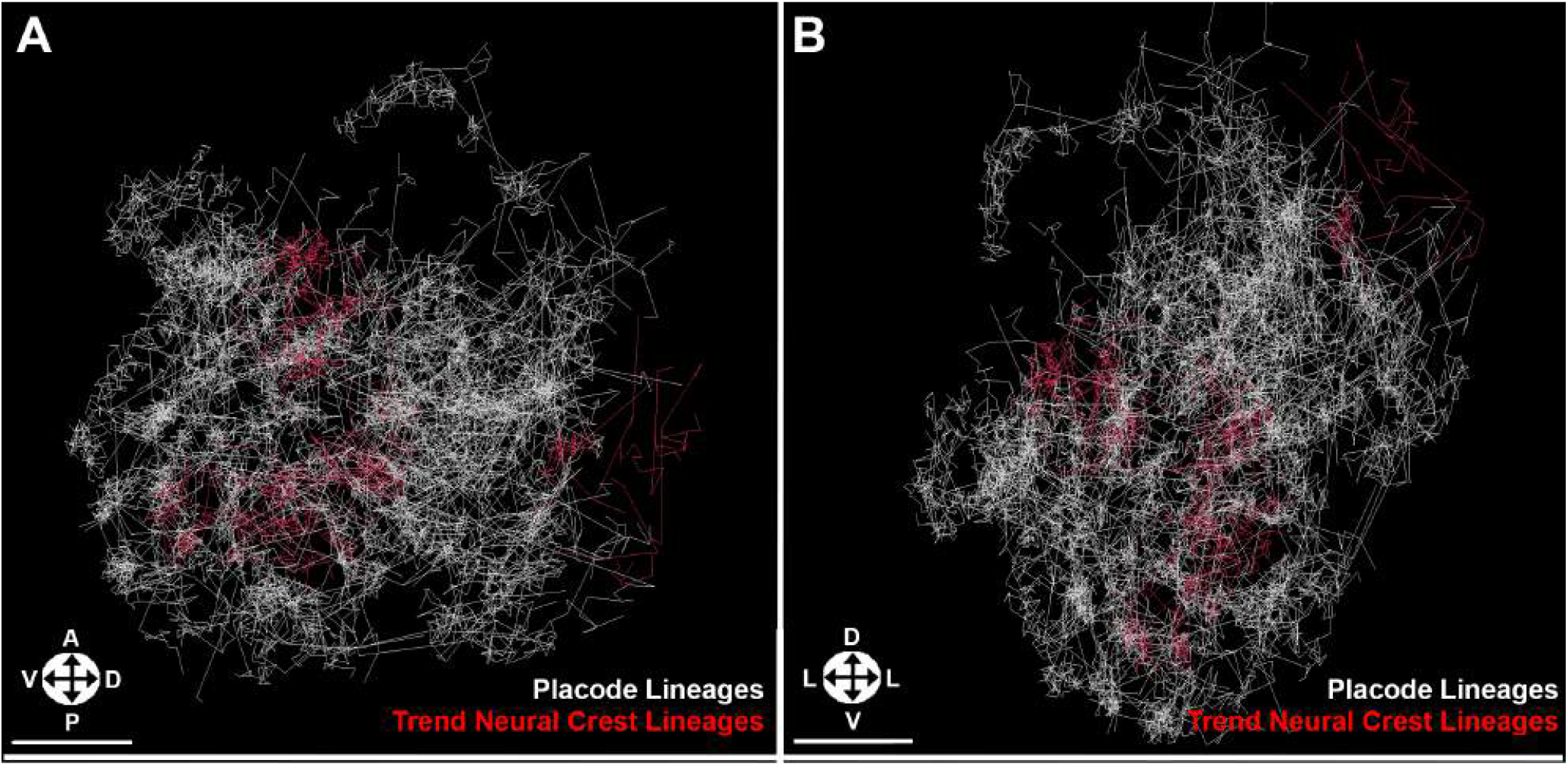
*Related to Figure 3*. Trend Neural Crest and Placode Lineages. (A) Lateral view of the olfactory epithelium showing trajectories of individual trend neural crest (red) and placode (white) lineages at 24-29.5 hpf. (B) 90 degree rotation of view shown in (A). Trend trajectories are not uniform in terms of spatiotemporal overlap with respect to placode trajectories. Orientation arrows: A, anterior; P, posterior; D, dorsal; V, ventral; L, lateral. Scale bars: 15 μm.

**Figure S2.**
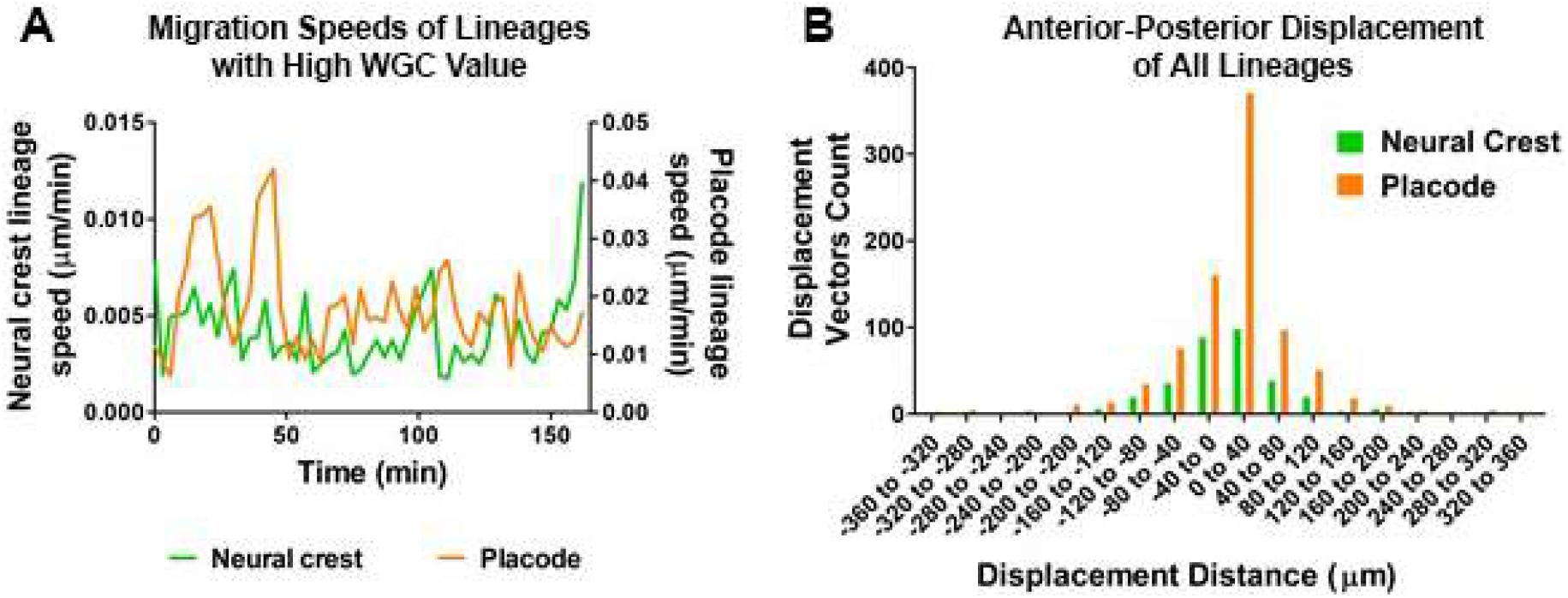
*Related to Figure 4*. Wiener-Granger Causal Analysis and Descriptive Statistics. (A) Speed of a representative pair of correlated lineages over time indicating harmonic oscillation. Average speed of an NC lineage (green, speed on the left y axis) and a placode lineage (orange, speed on the right y axis). These lineages are termed ‘correlated’ because they were related by Wiener-Granger causality at a high level of statistical significance (p<0.001). (B) Lineage displacement histogram presenting the distribution of averaged 3’ displacements over a 300’ window of observation (24.5-29.5 hpf) for NC (green) and placode (orange) lineages. Displacement vectors for each lineage have been sorted by bins of 40 μm. Negative values are anterior, positive values are posterior. The majority of values in the 0-40 μm bin for placodes is concentrated near a zero displacement, consistent with a relatively unbiased stochastic mechanism.

**Video S1. *Related to Figure 1*. Example of lattice light-sheet imaging of neural crest ingression into the olfactory epithelium.** Four different views are shown at nearly isotropic resolution of ~6.6 hours of olfactory development with 3’ time points. *Tg(OMP2k:lyn-mRFP* marks ciliated OSNs and *Tg(TRPC24.5k:gap-Venus)* marks microvillous OSNs.

**Video S2. *Related to Figures 2–5*. Algorithmically generated tracks of neural crest- and placode-derived cells during development of the olfactory epithelium.** Cellular nuclei, marked by *Tg(−8.4neurog1:nRFP)*, were used to determine positions of cells and tracks representing each cell’s motion. Shown and analyzed is a duration of 24-29.5 hpf.

**Video S3. *Related to Figure 3*.** Cellular trajectories of placode-derived cells (white) and NC-derived cells (blue) from lineages categorized in the volatility analysis as dispersed. Note the distinctive lack of overlap between placode- and NC-derived trajectories. Shown and analyzed is a duration of 24-29.5 hpf.

**Video S4. *Related to Figure S1*.** Cellular trajectories of placode-derived cells (white) and NC-derived cells (red) from lineages categorized in the volatility analysis as trend. Contrast with Video S3; unlike dispersed trajectories, trend trajectories do overlap with placode-derived trajectories. Shown and analyzed is a duration of 24-29.5 hpf.

**Supplemental Table 1:** Analytical Inventory

| **Technique** | **Purpose** | **Utility** | **Requirements** | **Findings** |
| --- | --- | --- | --- | --- |
| Exploratory Data Analysis | Illuminate statistical features of displacement data. | Inform model selection choice based through evidence indicating useful quantitative treatment of variables. | Acceptable optimization of trade-off between accuracy and precision. | Displacements are normally distributed, and speed variation is correlated between the neural crest and placode. |
| Algorithmic Pattern Classification | Assessment of extent to which computational statistical analysis of data can support a qualitative discrimination. | Provides support for the quantitative treatment of variables supported by exploratory data analysis. | Informed choice of a statistical pattern classification method. | Algorithmic discrimination between neural crest and placode cell identities performed much better with lineal than cellular data. |
| Volatility Analysis | Similar to standard deviation, except centered around a trend displacement. | Provides evidence of the dispersal of a set of sample displacements around a trend of motion. | Roughly normally distributed samples. | Neural crest-derived lineages clustered into two distinct groups, one highly volatile and one comparatively non-volatile. |
| Wiener-Granger Causal Analysis | Detection of predictive relations between trajectories. | Provides indirect evidence of mechanism coordinating motions of cells. | Linear dynamical mechanism of coordination. | Asymmetric relations of predictive information dependence between placode, NC 'trend' and NC 'dispersed' clusters of lineages. |
| Cointegration Test | Dynamically equilibrating vector error-correction model. | Provides evidence of the extent to which a pair of time series exhibit a pair-wise relationship that is subject to dynamical equilibration. | Frequently sampled measurements. | The motion of neural crest- and placode-derived lineages appears coordinated such that an average separation is maintained. Stochastic shocks are corrected as if by a spring mechanism. |

## Methods

### Zebrafish

Animals were treated and cared for in accordance with the National Institutes of Health Guide for the Care and Use of Laboratory animals and under the protocols of the Institutional Animal Care and Use Committee of HHMI Janelia Farm. Embryos were grown, staged, selected for transgenic markers, and treated to inhibit pigmentation as previously described(Kimmel et al., 1995; Rajan et al., 2018). Transgenic lines used were: *Tg (−4.9sox10:eGFP)* (Wada et al., 2005) = Sox10:eGFP; *Tg(−8.4neurog1:nRFP)* (Blader et al., 2003)= Ngn1:nRFP; *Tg(OMP2k:lyn-nRFP)/rw035* (Sato et al., 2005); *Tg(TRPC24.5k:gap-Venus)/rw037* (Sato et al., 2005). Zebrafish matings yielded compound heterozygote embryos for experiments.

### Live Imaging

Imaging preparation for confocal time-lapse imaging was done as previously described(Rajan et al., 2018). Imaging preparation for LLS imaging was done as previously described (Chen et al., 2014), with the following modifications: Embryos were embedded in 0.3% low-melting agarose (Sigma-Aldrich A4018) prepared in 30% Danieau solution with standard working concentrations of tricaine anesthetic and N-phenylthiourea (to prevent pigmentation) and suspended in 30% Danieau maintained at 28.5°C. Prior to and during experimentation, embryo staging was carefully monitored. Imaged embryos were compared to embryos embedded in .3% agarose but not imaged and to embryos that had remained in egg water. No discernable developmental delay or evidence of phototoxicity was observed in the olfactory organs of imaged embryos as compared to control groups (data not shown). Even in extreme cases of longer time-lapse imaging such as that shown here (Figure 1B) after 16 hours of continuous imaging with 3’ time points, only a slight truncation of the posterior tail region of the embryo was observed and was likely due to our mounting methodology that placed greater constraint upon that region as opposed to the anterior region of interest. Shorter time-lapses did not yield this truncation. Several hours after returning imaged embryos to egg water, imaged and control embryos were indistinguishable from each other and grew normally.

### Data Conversion and Analysis

Cell movements were tracked in Imaris Software (Bitplane, Inc.) using the Spots function. Raw tiff files were either analyzed directly or first registered and deconvoluted to correct for drift and background, respectively. Drift correction was implemented using Imaris 8.2.1. Tracking of cell trajectories and determination of lineal relationships was implemented using Imaris 8.3.1 through the ‘ImarisTrackLineage’ functionality. This construction of trajectories resulted in 250 cellular trajectories (63 NC-derived, 112 placode-derived, and 75 in the olfactory bulb) and 32 lineal trajectories in the OE (12 NC-derived and 20 placode-derived). There are two possible interpretations of individual lineage tracing results (Figure 2A). Lineages can be interpreted as traces of cell divisions or as spatial clustering of cell tracks that were within the lineage tracing algorithm’s threshold for lineal categorization. Either determination is compatible with our subsequent analysis of the dynamical patterning of cell migration and rearrangement. For effectively tracked cell divisions, our results provide evidence of lineally-specific dynamical patterning; otherwise, the results provide evidence of spatially-specific dynamical patterning.

### Analysis Methodology

Supplemental Table 1 provides a summary of employed econometric and finance-theoretic quantitative analysis methodology, including each technique’s purpose, utility, data requirements, and findings. The following sections discuss each method in detail.

### Principal Components Analysis and Histograms

A principal components analysis (PCA) generates a representation of the data in new coordinates, which are linear combinations of the original coordinates. These coordinates are uncorrelated and sequentially account for as much variance in the data as possible. We performed PCA on cellular (63 NC-derived and 112 placode-derived) and lineal (12 NC-derived and 20 placode-derived) trajectories using Matlab. Histograms of cellular and lineal distributions of displacement were constructed using R. Lineal distributions were constructed from individual time series by averaging the positions of all descendants of cells in existence at the beginning of imaging. Analyses of the distribution of cellular and lineal displacements along the three axes suggested that cellular trajectory data was less analytically revealing than lineal data (a Kolmogorov-Smirnov test, used to test for normality of a distribution, indicated normality of lineal, as opposed to cellular, displacement distributions; data not shown).

### Automated Cell Lineage Categorization

A maximum *a posteriori* pattern classifier was constructed based on training data associated with the 175 cellular and 32 lineal trajectories in the OE, with lineages manually identified as either NC- or placode-derived based on fluorescent markers. The classifier categorized lineages based on minimizing the Mahalanobis distance between measurements of *N* types to N-dimensional Gaussian distributions fit to each category based on training data. The categories considered were velocity and acceleration on each anatomical axis, start to finish distance and displacement, mean squared displacement, and coefficient of variation of velocity and acceleration. These calculations were implemented in Matlab. Optimal performance was attained when anteroposterior axial net displacement, anteroposterior axial average velocity, and dorsoventral axial average velocity were used as the measurement types.

### Wiener-Granger causality, Cointegration, and Volatility

Average lineal positions and velocities were calculated using Matlab scripts. Since we sought to statistically assay causal relations between cellular trajectories, we deployed a particular type of Wiener-Granger causality (WGC) test. Informally, WGC is said to exist between time series X and Y when an estimator predicts future values of X better when it is given the past values of X and the past values of Y than when it is given the past values of X alone. This mathematizes the intuitions that causes should precede effects and that knowledge of causal variables improves retrodictability of effectors (Granger, 1969). A significant difficulty in interpreting the results of WGC analysis in general is the problem of spurious correlation, i.e. it is possible for artefactual structures in data to lead to apparently statistically significant findings of WG causality despite the absence of an actual mechanism relating the structures. To rule out spurious correlations, we focused on a restricted version of WGC analysis by employing statistical tests of cointegration. Cointegration tests were deemed appropriate given the approximate normality of lineal anteroposterior displacement distributions. This was determined through construction of histograms with bin width of 40 μm (Figure S3B). The normality of this distribution indicated the plausibility of a model based on a stochastic ordinary differential equation model that couples a deterministic process to a stochastic process. In the case of cointegration, the deterministic process is the vector error-correction function acting on anteroposterior displacements, while the stochastic process is the source of random perturbations to the equilibrium trend. More formally, wide-sense stationary time series of order N for which a (wide-sense) stationary linear combination exists of order strictly less than N are termed cointegrated. Informally, time series are cointegrated when their fluctuations are consistent with the action of either a common trend or an equilibrating, spring-like mechanism acting on both of them to maintain a common separation or “spread” (this result is known as Wold’s Theorem, (Wold, 1954)).

Cointegration relations were computed between 34 time series representing the average position of a given cell lineage (12 epithelium-directed NC; 2 bulb-directed NC; 20 epithelial placode), 10 individual cell trajectories of non-migrating NCCs in the cavity, and 66 individual motile cell trajectories (20 epithelium-directed NC; 4 bulb-directed NC; 42 epithelial placode; not all of the 175 cellular trajectories in the OE were subject to cointegration analysis because some did not have continuous data during the time window of analysis or exhibited tracking errors that were qualitatively discernible).

Cointegration relations were analyzed using Johansen’s statistical test (Johansen, 1991) as implemented in the freely available R package *urca* (maintained by Bernhard Pfaff). Heat maps were constructed in MatLab.

Volatility was deployed as a statistical measure of the dispersion of displacements about a displacement trend. The calculation is identical to a standard deviation calculation, except the residuals are computed with respect to the displacement trend rather than the displacement mean. Volatility was calculated using Matlab scripts.

### Cumulative Work Calculation

The *urca* package provided half-life estimates for each pairwise cointegration relation. These half-lives were converted into spring coefficients associated with the interpretation of cointegration as a first-order linear equilibrating mechanism subject to stochastic perturbations. The first-order models were treated as representations of hypothetical elastic tethers acting on lineal trajectories, with the average separation between pairs of lineages interpreted as the equilibrium separation associated with the elastic tether. We restricted our attention to placode-NC lineage pairs, since these appeared highly statistically significant and indicated a previously undescribed modality of interaction between two subpopulations. The anterior-posterior displacement of each NC lineage was treated as an oriented distance. The dot product of this oriented distance and the force associated with separation from equilibrium for each pairing of a given NC lineage with each placode lineage was calculated, and the total was interpreted as a proxy for the net biophysical work done by the hypothetical elastic tether.

